# A Size Filter Regulates Apical Protein Sorting

**DOI:** 10.1101/2023.07.13.548868

**Authors:** Christian de Caestecker, Ian G. Macara

## Abstract

Despite decades of research, apical sorting of epithelial membrane proteins remains incompletely understood. We noted that apical cytoplasmic domains are smaller than those of basolateral proteins; however, the reason for this discrepancy is unknown. We investigated whether a size barrier at the trans-Golgi network (TGN) might hinder apical sorting of proteins with large cytoplasmic tails. We focused on Crb3 and Ace2 as representative apical proteins with short cytoplasmic tails. By incorporating a streptavidin-binding peptide, these proteins can be trapped in the endoplasmic reticulum (ER) until addition of biotin, which triggers synchronous release to the Golgi and subsequent transport to the apical cortex. Strikingly, departure from the Golgi could be significantly delayed simply by increasing cytoplasmic bulk. Moreover, large and small Crb3 segregated into spatially distinct Golgi regions as detected by super resolution imaging. Biologically, Crb3 forms a complex through its cytoplasmic tail with the Pals1 protein, which could also delay departure, but although associated at the ER and Golgi, we found that Pals1 disassociates prior to Crb3 departure. Notably, a non-dissociable mutant Pals1 hampers the exit of Crb3. We conclude that an unexpected mechanism involving a size filter at the TGN facilitates apical sorting of proteins with small cytoplasmic domains and that timely release of Pals1, to reduce cytoplasmic domain size, is essential for the normal kinetics of Crb3 sorting.

## Introduction

An essential aspect of epithelial cell biology is the polarized sorting of membrane proteins to the basolateral and apical plasma membranes^1,2^. This differential sorting, which occurs at the trans-Golgi network (TGN), maintains the identities and functions of these two cortical regions, and is disrupted in multiple human diseases^2^. Short zip codes within the sequences of basolateral membrane proteins provide sorting information, but apical sorting mechanisms are still not fully understood^3,4^. Clustering through lipid phase separation or pH-dependent protein-glycan interactions, and indirect delivery via transcytosis, have been implicated in specific cases but are not universally applicable^5-9^. An intriguing feature of many apical membrane proteins is their short or non-existent cytoplasmic domains. Some apical proteins are GPI-linked and do not penetrate through the plasma membrane to the cytoplasmic side^10^. Others have short cytoplasmic tails^11^. In contrast, basolateral plasma membrane proteins often have large cytoplasmic domains that frequently form complexes with other components. Examples include receptor tyrosine kinases and cadherins^12,13^. The reasons for this apparent structural asymmetry between membrane proteins at the apical versus basolateral regions of the plasma membrane are unknown. We wondered, however, if this asymmetry might be connected to differential sorting, and considered the possibility that a diffusion barrier at the TGN might hinder accumulation of certain cargoes with large cytoplasmic domains into apical carriers, by analogy with the nuclear pore barrier. Diffusion of membrane proteins across the nuclear pore to access the inner nuclear envelope is only possible for proteins with small cytoplasmic domains because of steric hindrance^14,15^. Membrane proteins with large cytoplasmic domains must use an energy-dependent mechanism that requires karyopherin transporters. Primary cilia also maintain a diffusion barrier at the transition zone^16^.

Studying anterograde traffic of proteins to the plasma membrane poses a challenge, as negligible amounts are present in the biosynthetic route at steady state. Even when intracellular trafficking can be visualized at steady state, it is challenging to discriminate between anterograde and retrograde trafficking events. To circumvent these problems, we employed the Retention Using Selective Hooks (RUSH) system to enable synchronized anterograde transit of specific proteins^17^.

To test the size filter concept, we used the apical transmembrane polarity protein Crumbs3, which has a cytoplasmic C-terminal tail consisting of only 39 amino acid residues^11^. An SBP-Halotag or -GFP attachment at the N-terminus enabled visualization and synchronous release from the endoplasmic reticulum (ER) upon biotin addition. To increase cytoplasmic domain size in a scalable fashion, we used a chemical dimerizer system. This system demonstrated a significant impairment of TGN exit when one or more SNAPtags or mCherry were attached by addition of dimerizer. Interestingly, Crb3 associates with a large adapter protein, Pals1, which would be predicted to impair TGN export when bound^18,19^. We discovered, however, that Pals1 disassociates from Crb3 prior to Crb3 accumulation in tubules and exit from the TGN. A deletion mutant of Pals1 that remains bound to Crb3 substantially delays exit. These data suggest that the release of Pals1 reduces the cytoplasmic footprint of Crb3, ensuring its timely release through the size filter from the Golgi to the apical plasma membrane.

## Results

### A survey of apical versus basolateral membrane proteins reveals a significant cytoplasmic domain size difference

To determine if the anecdotal indication of a size differential between apical and basolateral protein cytoplasmic domains is real, we surveyed all the epithelial membrane proteins in the PolarProtDB database^20^. Entries were annotated by cytoplasmic domain length. For multi-pass proteins the longest cytoplasmic segment was chosen. Noncovalent associations with other proteins were ignored. Data are presented in **Fig. 1** and **Auxiliary Material 1**, where a clear bias is apparent towards smaller cytoplasmic domains for apical versus basolateral membrane proteins (**Fig. 1A**) while there is a much smaller difference in total length (**Fig. 1C**). The survey is dominated by the very large number of multipass solute carriers that are distributed about evenly between the basolateral and apical domains. Excluding this group increases the size differential between apical and basolateral cytoplasmic sequences, again with little difference in total length (**Figs. 1B, D**). The reason for such a discrepancy has not previously been addressed. We reasoned, however, that perhaps a size filter or diffusion barrier in the TGN preferentially enables such proteins to be sorted into apical carriers.

**Figure 1.**
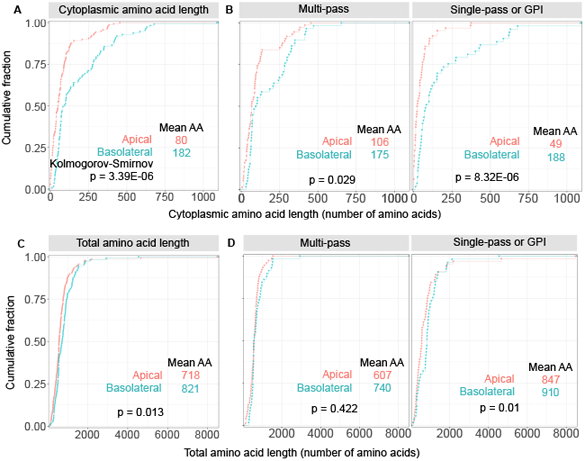
A survey of apical versus basolateral membrane proteins reveals a significant cytoplasmic size domain difference. (**A**) Cumulative distribution function of the cytoplasmic amino acid lengths for polarized membrane proteins stratified by apical versus basolateral localization based on localizations listed in *Zeke et al 2021*. (**B**) The same dataset was also stratified to compare multi-pass versus single-pass/GPI-linked proteins. A two-sided Kolmogorov-Smirnov test was used to assess the statistical significance of discrepancy between cumulative distributions. Apical n = 138 proteins; apical multi-pass n = 74; apical single-pass or GPI-linked n = 64. Basolateral n = 111 proteins; basolateral multi-pass n = 58; basolateral single-pass n = 53. (**C**) Reference cumulative distribution function of the total amino acid lengths for all apical versus basolateral membrane proteins from the same dataset, or (**D**) stratified for multi-pass versus single-pass proteins.

### Synchronizing anterograde traffic of the apical protein Crumbs3

To study anterograde traffic of apical proteins, we implemented the Retention Using Selective Hooks (RUSH) system for use with the apical membrane protein Crumbs3 (Crb3)^17^. The RUSH system retains a protein of interest at the endoplasmic reticulum (ER) by fusing it to a streptavidin-binding peptide (SBP), which interacts with streptavidin fused to a KDEL ER-localization sequence (StrKDEL) (**Figs. 2A, B**). The SBP-StrKDEL interaction is disrupted upon addition of biotin to the culture medium, enabling synchronized release of the SBP-fusion protein from the ER to resume its normal trafficking itinerary. By expressing SBP-Halotag-Crb3 and StrKDEL in live polarized Eph4 mammary epithelial cells, we can follow anterograde transit of Crb3 to the apical surface using confocal microscopy. Prior to biotin addition, SBP-Halo-Crb3 was absent from the apical surface and cellular junctions and was instead retained at the ER. Biotin addition triggered SBP-Halo-Crb3 release from the ER and synchronous transfer to the Golgi apparatus (GA), before being delivered to the apical surface over the course of 2 hrs (**Figs. 2C, Extended Data Fig. 1A, and Movie 1**). Quantitative analysis at single cell resolution was performed as described in the Methods, to calculate the Golgi dwell time. Additionally, we created a knock-in Eph4 cell line by CRISPR/Cas9-mediated gene editing, in which the Crb3 locus was modified to add SBP-Halotag at the N-terminus following the signal peptide (**Extended Data Fig. 1 B**,**C**). Endoge-nous Crb3 is expressed at very low levels in Eph4 cells but at steady state the Halo-Crb3 was detectable at cell junctions (**Extended Data Fig. 1D**). The cells were also engineered to express StrKDEL and in the absence of biotin the Halo-Crb3 was absent from cell junctions but detectable as intracellular puncta (**Extended Data Fig. 1E**). Importantly, upon release from the ER, the Halo-Crb3 accumulated in the Golgi and later at the plasma membrane, with similar kinetics to the over-expression system (**Fig. 2D, and Movie 2**). These data argue that the sorting itinerary is not impacted by accumulation of exogenously expressed Crb3 in the ER, and that the system provides a realistic measure of transit rates.

**Figure 2.**
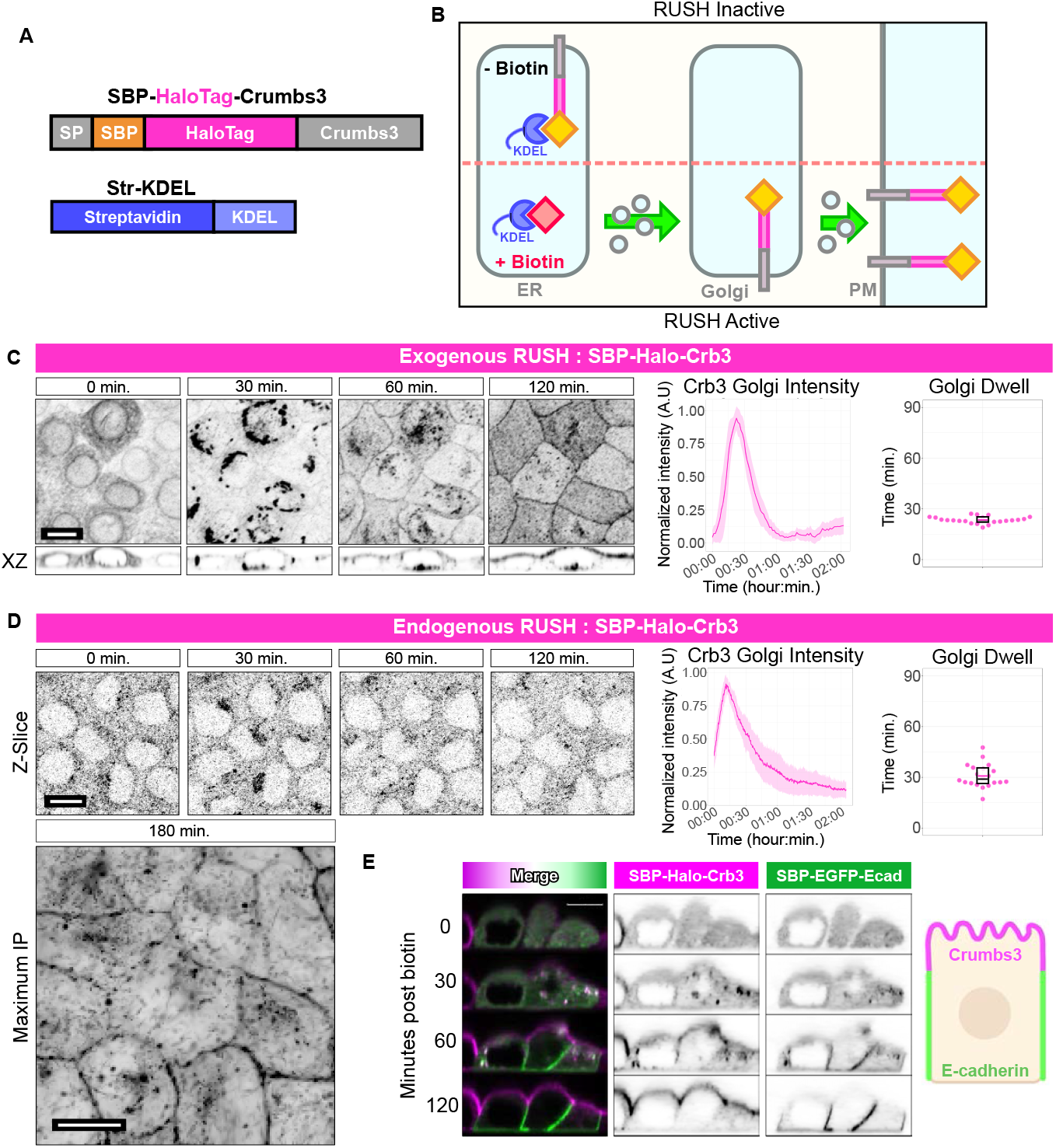
Synchronizing anterograde traffic of the apical protein Crumbs3. **(A)** Schematic of SBP-Halo-Crb3 construct. SP = Signal Peptide. SBP = Streptavidin Binding Peptide. **(B)** Schematic of RUSH system. In the absence of biotin, SBP-tagged cargo is retained in the ER through interaction with ER-localized Streptavidin-KDEL. Addition of biotin to culture media disrupts interaction enabling forward trafficking of SBP-fused cargo. **(C)** Confocal imaging of exogenous SBP-Halo-Crb3 RUSH in polarized Eph4 cells at intervals following addition of biotin to culture medium. Maximum intensity projections or XZ orthogonal views. Scalebar 10 μm. Middle panel: Aggregate quantification of Crb3 Golgi intensity during RUSH experiment. Solid line = mean; ribbon = SD. Right panel: Crb3 Golgi dwell time between prior and post 50% quantiles. n = 22 cells. **(D)** Same as **C**, but for endogenously labelled SBP-Halo-Crb3. Images acquired as single Z-slice, with a full Z-stack acquired at conclusion of the assay at t = 180 min. Scalebar 10 μm. n = 22 cells. **(E)** Confocal XZ orthogonal views of simultaneous exogenous RUSH SBP-Halo-Crb3 (magenta) and SBP-EGFP-Ecadherin (green) in polarized Eph4 cells at intervals following addition of biotin. Scalebar 10 μm.

To ensure that cargo-specific itineraries are maintained, we performed a multi-cargo RUSH experiment using both apically destined Crb3 and laterally destined E-cadherin RUSH constructs within the same cells. These cargos were delivered independently to their appropriate membranes (**Fig. 2E and Movie 3**). Importantly, the presence of the E-cadherin RUSH construct did not impede Crb3 apical delivery, suggesting that the biotin-triggered bolus of membrane proteins does not saturate the ER-Golgi system.

### Increasing the cytoplasmic domain size of Crb3 impedes anterograde trafficking

To assess the effect of cytoplasmic domain size on sorting dynamics of apical proteins, we implemented the FKBP-FRB chemical-inducible dimerization (CID) strategy to enable recruitment of varioussized FRB-tagged cargoes to the intracellular domain of SBP-Halo-Crb3-KBP by addition of A/C Heterodimerizer (**Fig. 3A, B**). First, we assessed the localization of SBP-Halo-Crb3-FKBP versus SBP-EGFP-Crb3 in polarized Eph4 cells at steady state, to ensure that fusing FKBP to the cytoplasmic face did not impair apical localization (**Fig. 3C**). Next, we introduced the StrKDEL ER hook to enable use of these constructs with the RUSH system, into cells that express both SBP-Halo-Crb3-FKBP, which can dimerize with FRB-tagged cargo in the cytosol, as well as SBP-EGFP-Crb3, which cannot dimerize and serves as an internal control. The internal control SBP-EGFP-Crb3 has identical trafficking dynamics to SBP-Halo-Crb3 (**Extended Data Fig. 2A**). In addition, we expressed recruitable cargos of various sizes in stoichiometric excess: SNAPx1-FRB (287 AA), SNAPx2-FRB (478 AA), or SNAPx3-FRB (669 AA) (**Fig. 3A, B**). The SNAPtag was chosen because it is monomeric and can be labeled with fluorescent dyes. When dimerizer is added to these cells, SNAP-FRB colocalizes with SBP-Halo-Crb3-FKBP at the ER (**Extended Data Fig. 2B**). In the absence of dimerizer, the kinetics of Golgi entry and departure for FKBP-fused Crb3 and control RUSH constructs were identical (**Fig. 3D-F and Movie 4**). However, when cells were treated with dimerizer prior to the RUSH assay, the recruitment of SNAP-FRB cargo to Crb3-FKBP substantially lengthened transit times (**Figs. 3D-F and Movie 4**). Surprisingly, however, transit duration did not scale with increasing number of SNAP-tag moieties on the recruited FRB cargo, suggesting the traffic delay is governed by a defined cytoplasmic size threshold (**Figs. 3G, Extended Data Fig. 2C**).

**Figure 3.**
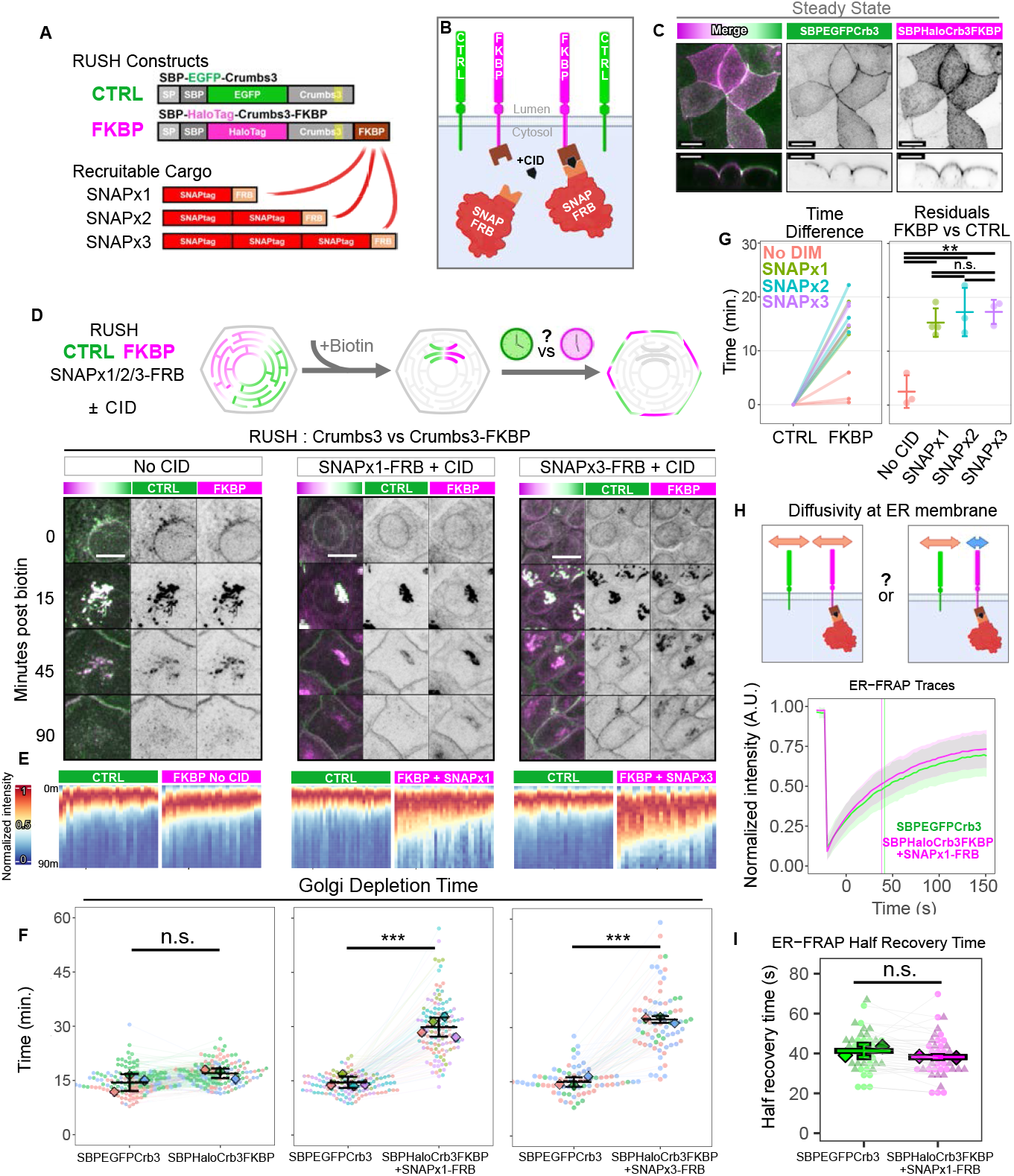
Cytoplasmic domain size regulates dynamics of apical trafficking. **(A)** Construct designs. SBP-EGFP-Crb3 serves as an internal control. SBP-Halo-Crb3-FKBP, which can be induced to dimerize with FRB-labelled cargo in the presence of a chemical inducible heterodimerizer (CID). Recruitable cargos are SNAP-FRB constructs with 1 - 3 SNAPtag moieties. **(B)** Schematic of constructs in the cell. **(C)** Steady-state localization of SBP-EGFP-Crb3 and SBP-Halo-Crb3-FKBP in live polarized EpH4 cells. Confocal maximum intensity projection. Orthogonal XZ views are denoised and visualized at a plane indicated by the arrowhead. Scalebar 10 μm. **(D)** Experiment schematic and still images of control SBP-EGFP-Crb3 (CTRL,green) and SBP-Halo-Crb3-FKBP (FKBP,magenta) during RUSH experiment in presence of SNAPx1-FRB and vehicle, or SNAPx1-FRB and CID, or SNAPx3-FRB and CID (left to right). Scalebar 10 μm. **(E)** Heatmaps of fluorescence intensity at the Golgi of SBP-EGFP-Crb3 or SBP-Halo-Crb3-FKBP in the prescence of SNAP-FRB ± CID. Each column represents channel normalized Golgi intensity over time for a single cell in the above experiment. **(F)** Aggregated Golgi depletion time data. Each point is a cell, and each line joins paired measurements within the same cell. Datapoints are colored by experiment, with diamonds representing the mean of each experiment. n *≥* 74 total cells and N *≥* 3 experiments per condition. Student’s t-test. n.s. p = 0.180; ***p = 5.52e-05; ***p = 4.88e-05. **(G)** Golgi depletion time residuals comparing FKBP versus control from **D**. One-way ANOVA + Tukey’s HSD. n.s. p > 0.841; **p < 0.0022. **(H)** Diffusion schematic and aggregate FRAP traces of ER-localized SBP-EGFP-Crb3 (green) or SBP-Halo-Crb3-FKBP (magenta) within the same cell in the presence of SNAPx1-FRB and CID. Solid line: mean. Ribbon: SD. **(I)** Half-maximum recovery times from **H**. Each point is a cell, and each line joins paired measurements within the same cell. n = 42 total cells, N = 2 experiments. Student’s t-test n.s. p = 0.164.

To test if the SNAPtag itself might impact the kinetics of Crb3 traffic through the Golgi, we replaced it with a mCherry-FRB fusion construct. Importantly, dimerization of this protein to the Crb3-FKBP also significantly inhibited anterograde traffic (**Extended Data Fig. 2D**), demonstrating that the effect is independent of the nature of the cytoplasmic attachment.

Another possible explanation we ruled out was that bulky FKBP-FRB groups block recruitment of AP trafficking adaptors to Crb3. AP-2 binds to a region overlapping the C-terminal PDZ-binding motif of Drosophila Crb and is responsible for its endocytosis^21^. The exocytic adaptor AP-1 shares a similar recognition sequence. However, the dynamics of a Crb3 mutant lacking this domain were identical to the full-length protein (**Extended Data Fig. 2E**).

We next considered the possibility that the reduced transit rates for Crb3 with attached SNAPtags or mCherry might be a consequence of reduced intrinsic diffusion rates. However, Houser and colleagues observed that the size of extramembrane domains has no influence on intrinsic 2D diffusivity in membranes^22^. Moreover, the diffusivities of membrane proteins were 1-2 orders of magnitude below those expected for 3D diffusion of similar-sized globular proteins in solution, suggesting that viscous drag within the membrane is dominant over hydrodynamic drag on the cytoplasmic do-mains of the proteins^23^. Additionally, molecular crowding in the membrane inhibits diffusivity by steric exclusion independently of protein size^22^. Consistent with these results, we detected no significant difference experimentally in the rates of recovery after photobleaching for Crb3-FKBP at the ER with SNAP-FRB attached when compared with the short form Crb3 (**Fig. 3H, I**). We therefore conclude that diffusion rates cannot explain the effect of increased cytoplasmic domain size on apical membrane protein sorting in the absence of a barrier.

### Apical proteins with small cytoplasmic domains are preferentially sorted into a distinct region of the Golgi

One key prediction of the size filter hypothesis is that such a filter would gate access to a sub compartment of the TGN, from which apically targeted cargos with bulky cytoplasmic domains would be excluded. To test this prediction, we repeated the Crb3 RUSH experiments ± dimerizer to manipulate the size of the cytoplasmic domain and visualized the cells live, using super resolution Airyscan confocal microscopy. We sought to resolve sub-compartments that would contain the GFP Crb3 but not the bulkier Halo-Crb3-FKBP linked to a SNAPx1-FRB. As shown in **Fig. 4A-C**, distinct areas of the Golgi contain the GFP-Crb3 but not the dimerized Halotag version. Together, these data support the concept of a cytoplasmic domain size-dependent sorting mechanism at the Golgi for apically destined membrane proteins.

**Figure 4.**
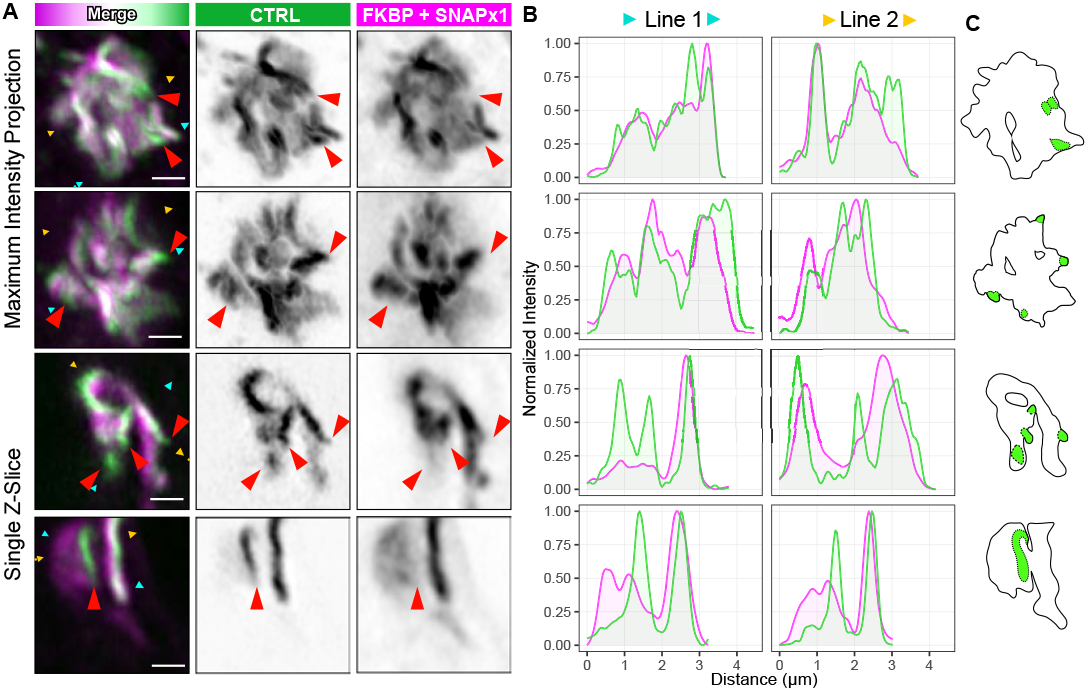
Apical proteins with small cytoplasmic domains are preferentially sorted into a distinct region of the Golgi. **(A)** Representative Zeiss Airyscan confocal images of the Golgi during RUSH in cells expressing SBP-EGFP-Crb3 (green) and SBP-Halo-Crb3-FKBP (magenta) coupled to SNAPx1-FRB via CID, captured 20 minutes after biotin addition. Red arrowheads indicate examples of regions enriched for SBP-EGFP-Crb3. Scalebar 1 μm. **(B)** Line scan quantification for **A**. Plots are normalized intensity values of an invisible line connecting cyan or yellow lines in **A. (C)** Cartoon depiction of SBP-EGFP-Crb3 enriched regions.

### The Ace2 apical membrane protein shows the same dependence as Crb3 on cytoplasmic domain size

A key consideration is whether the effects of cytoplasmic domain size on sorting dynamics are specific only to Crb3 or are generalizable to other apical transmembrane proteins. To address this point, we employed the RUSH-FKBP system using another, unrelated cargo, Ace2, which localizes to the apical surface in multiple epithelial cell types^24^. SBP was attached to the N-terminus of Ace2 after a signal peptide (**Fig. 5A**). Two RUSH versions were created as had been done with Crb3, one containing a GFP (mNeon) as a control and the other containing a Halotag plus a C-terminal FKBP for dimerization to SNAP-FRB, in addition to the SBP domain. Both versions localized exclusively to the apical surface of Eph4 epithelial cells in the absence of Streptavidin-KDEL (**Fig. 5B**). They also localized efficiently to the ER in cells containing the KDEL hook and were released by addition of biotin. The kinetics of delivery to and departure from the Golgi were faster than those for Crb3, but there was no difference in departure times between the mNeon control and the Halo-FKBP constructs. Strikingly, however, attachment of SNAP-FRB by addition of dimerizer significantly delayed TGN exit of the SBP-Halo-Ace2-FKBP. (**Fig. 5C-E, Extended Data Fig. 2E, and Movie 5**). These data demonstrate that the effect of increasing cytoplasmic domain size on apical sorting and transport is not restricted to Crb3.

**Figure 5.**
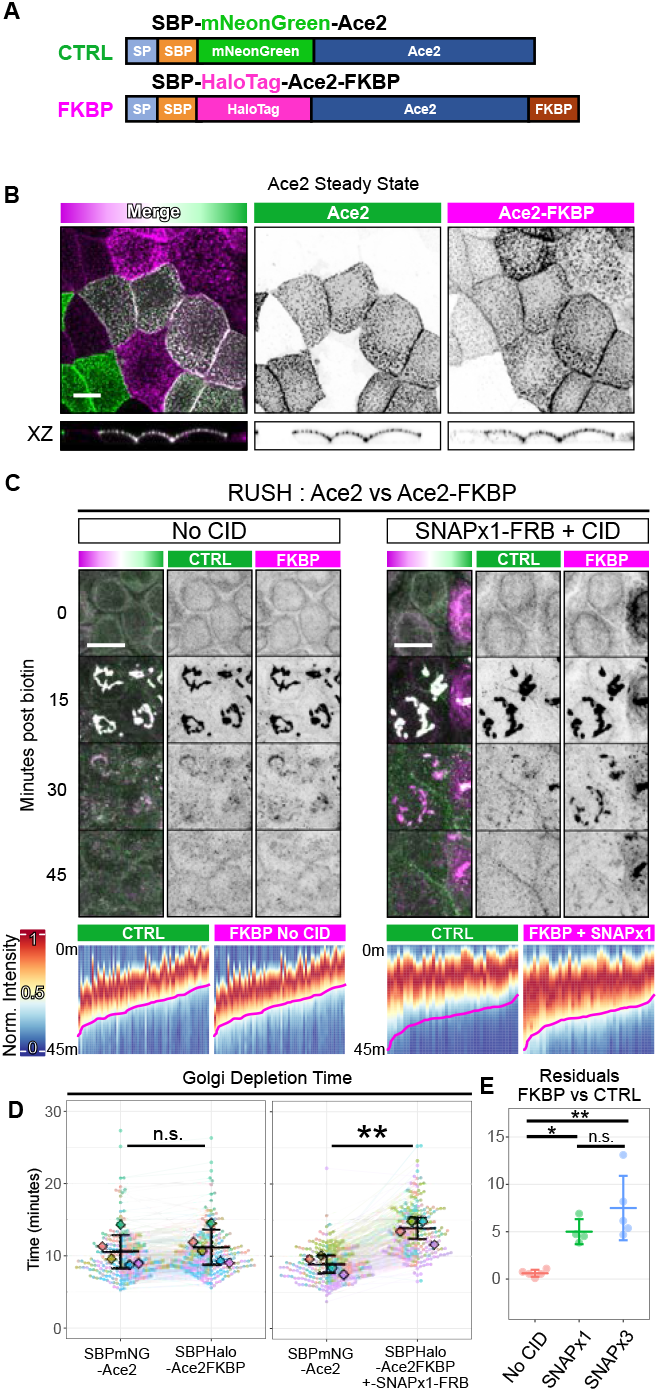
Ace2 sorting kinetics at the Golgi. **(A)** Schematic of Ace2 RUSH and dimerizer constructs. **(B)** confocal maximum intensity projections or XZ orthogonal views of Ace2 constructs in Eph4 cells lacking the ER hook, showing that both constructs are exclusively apical at steady state. Scalebar 10 μm. **(C)** RUSH kinetics of Ace2 constructs in the absence or presence of SNAP-FRB attached to the FKBP moiety at the C-terminus of the Halo-Ace2 construct. Top shows example images from the time courses. Confocal maximum intensity projection. Scalebar 10 μm. Bottom shows heatmaps of fluorescence intensity at the Golgi. Each column represents channel normalized Golgi intensity over time for a single cell in the above experiment. **(D)** Aggregated Golgi depletion time data. Each point is a cell, and each line joins paired measurements within the same cell. Datapoints are colored by experiment, with diamonds representing the mean of each experiment. n *≥* 183 total cells, N *≥* 4 experiments per condition. Student’s t-test. n.s. p = 0.697; **p = 0.00196 **(E)** Golgi depletion time residuals comparing FKBP versus control from D. One-way ANOVA + Tukey’s HSD. n.s. p = 0.245; *p = 0.029; **p = 0.001.

### Crb3 and Pals1 co-traffic from the ER to Golgi but dissociate prior to Golgi exit

Many membrane proteins have binding partners that would substantially increase their effective cytoplasmic domain sizes and would impede apical protein delivery if the associations occur in the ER or Golgi. To address this issue, we fixed SBP-Halo-Crb3 RUSH cells at a time when the Crb3 had accumulated at the Golgi, and immunostained a panel of proteins known to interact with Crb3 at the apical surface. Surprisingly, the Crumbs complex components Pals1 and Patj accumulated with Crb3 at the Golgi, but neither the tight junction protein ZO1, nor polarity proteins Par6 and aPKC colocalized with Golgi Crb3 (**Fig. 6A, Extended Data Fig. 3A**). To probe the biological implications of such interactions we studied Pals1, a cytoplasmic polarity protein of 74 kDa that binds to the C-terminal PDZ ligand sequence, E-R-L-I, on Crb3^25, 26^. Knockout of Pals1 did not affect the dynamics of Crb3 transit from the ER to PM, showing that the interaction is not essential for Crb3 anterograde transport (**Extended Data Fig. 3B**). Moreover, deletion of the C-terminal sequence to prevent Pals1 binding also did not prevent apical delivery of Crb3 (**Extended Data Fig. 1C**). These data raised the possibility that Pals1 protein associates with Crb3 only after exit from the TGN or only after delivery of Crb3 to the plasma membrane.

**Figure 6.**
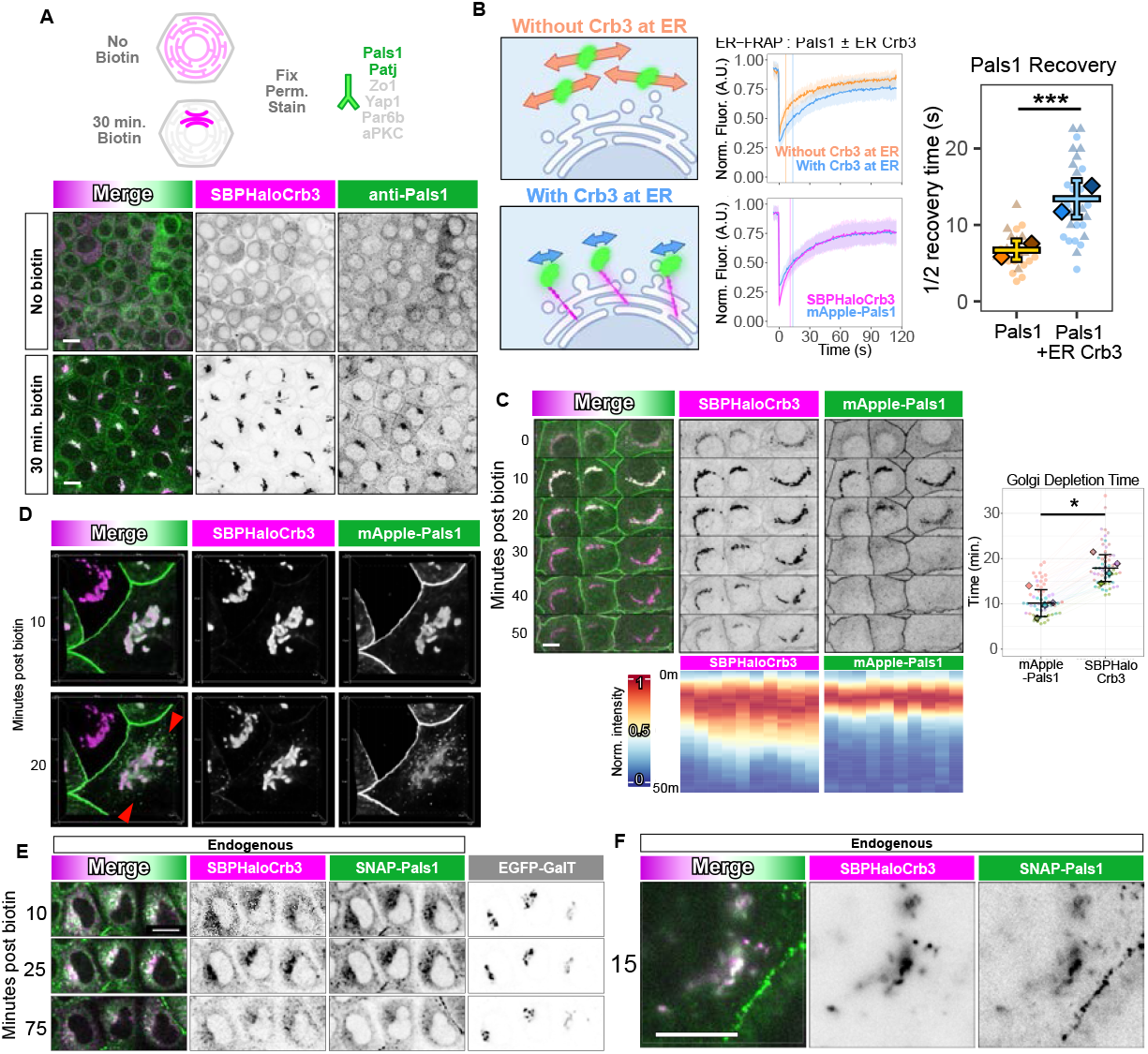
Pals1 dissociates from Crb3 at the Golgi just prior to Crb3 departure. **(A)** Images of SBP-Halo-Crb3 and endogenous Pals1 immunofluorescence staining in cells fixed before or after release from the ER by biotin addition. **(B)** FRAP analysis of SBP-Halo-Crb3 and mApple-Pals1 in the absence of biotin. Diagrams show predicted results assuming either that the two proteins are not associated at the ER or are associated. FRAP traces and recovery times support the hypothesis that Pals1 is bound to Crb3. n *≥* 25 total cells per condition from two experiments. Welch’s t test. ***p = 1.26e-07. **(C)** RUSH experiment showing that mApple-Pals1 departs the Golgi prior to SBP-Halo-Crb3, and Golgi depletion time quantification. n = 53 total cells. N = 4 experiments. Student’s t-test. *p = 0.010. **(D)** Confocal imaging detects mApple-positive, Halotag-negative puncta leaving the Golgi (red arrowheads), suggesting that the Pals1 is associated with vesicles. Confocal 3D projection. Bounding box: 30×30 μm **(E)** Confocal image slices of endogenously labeled SBP-Halo-Crb3 (JF646) and endogenously labeled SNAP-Pals1 (JF552) during RUSH. **(F)** Same as **E**, using near-TIRF, showing SNAP-Pals1 at both the Golgi and plasma membrane. All scalebars 10 μm.

To address this issue, we expressed an mApple-tagged Pals1 in the Crb3 RUSH competent Eph4 cells and tracked both proteins, before and after biotin ad-dition to release Crb3 from the ER. We determined by FRAP that mApple-Pals1 mobility was significantly reduced when Crb3 was retained at the ER, suggesting formation of a Crb3-Pals1 complex (**Fig. 6B**). Further, the recovery rates of mApple-Pals1 and SBP-Halotag-Crb3 were identical, also suggesting that Pals1 is associated with Crb3 at the ER (**Fig. 6B**).

Why then does Pals1 not inhibit exit of Crb3 from the TGN? When we activated RUSH, Crb3 and Pals1 accumulated at the Golgi apparatus synchronously (**Fig. 6C**). Surprisingly however, as traffic proceeded, Pals1 departed from the Golgi 7 min prior to Crb3 (**Fig. 6C and Movie 6**). Moreover, Pals1, which is a cytoplasmic protein, left the Golgi in the form of puncta (**Fig. 6D and Movie 7**). We have been unable so far to determine if these puncta are vesicles, or what if any additional proteins are interacting with Pals1 in them. To confirm that interaction at the Golgi was not an artefact of over-expression, we created a double homozygous knock-in Eph4 cell line to add an SBP-HaloTag to Crb3 and a SNAPtag to Pals1 at their endogenous loci (**Extended Data Fig. 3C-E**). Using these cells, were able to confirm co-trafficking between endogenous Crb3 and Pals1 (**Fig. 6E, F**). The low expression level of each protein prevented determination of when Pals1 departed the TGN. Nonetheless, we can conclude that Pals1 disassociation from Crb3 at the Golgi, rather than complex formation, is important for normal dynamics of Crb3 transit.

### Dissociation of Pals1 is essential for timely exit of Crb3 from the Golgi

Finally, we asked if dissociation of Pals1 is essential for rapid exit of Crb3 from the TGN. A truncating mutant of Pals1 (EGFP-Pals1ΔN) was generated that lacks the N-terminal portion required for Par6/Patj/Lin7c inter-actions, leaving intact the PDZ-SH3-GUK domains that are essential for binding to Crb3 (**Fig. 7A**). In Crb3-RUSH experiments, when EGFP-Pals1ΔN is expressed, we noted that it co-traffics with Crb3 but does not dis-sociate at the Golgi, as occurs with full length Pals1 (**Fig. 7B**). Consistent with our predictions, the transit time of Crb3 through the Golgi was dramatically lengthened when Pals1ΔN was expressed (**Fig. 7C**). We conclude that dissociation of Pals1 from Crb3 at the Golgi is an important regulatory step in the forward trafficking of Crb3 and delivery to the apical PM.

**Figure 7.**
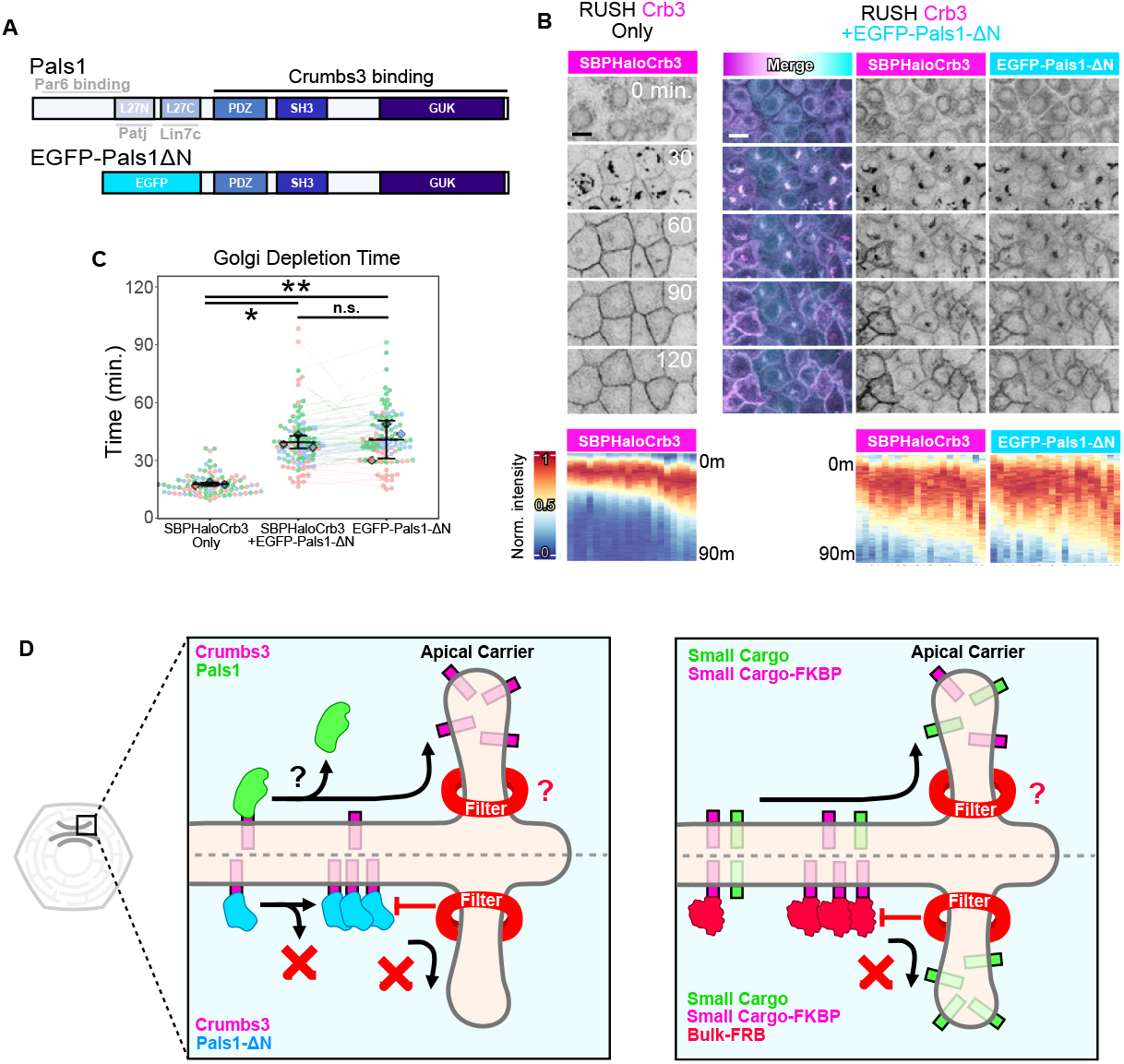
Dissociation of Pals1 is essential for timely exit of Crb3 from the Golgi. **(A)** Schematic of Pals1 domain structure, showing regions that bind Crb3 and other polarity proteins, and a mutant Pals1 (Pals1ΔN) that lacks the N-terminal domains and does not disassociate from Crb3 at the Golgi. **(B)** Confocal stills of SBPHalo-Crb3 release from the Golgi in the presence or absence of the Pals1ΔN mutant. Scalebars 10 μm. **(C)** Quantification of Golgi depletion kinetics from **B**. n *≥* 85 total cells per condition; N = 3 experiments. One-way ANOVA + Tukey’s HSD. n.s. p = 0.956; *p = 0.0106; **p = 0.0079. **(D)** Model of Golgi size filter. Left: Pals1 cotraffics with Crb3, but dissociates from it at the Golgi, reducing the cytoplasmic footprint of Crumbs3. This enables Crb3 to move past the size filter to enable sorting and exit from the TGN. Pals1ΔN is unable to dissociate from Crb3 at the Golgi, and therefore the Crb3-Pals1 complex cannot pass the size filter, hindering its ability to exit. Right: More generally, cargoes with bulky cytoplasmic footprints are occluded from apical sorting compartments at the TGN, but cargoes with smaller footprints are not, enabling entry and sorting.

## Discussion

How cells sort and deliver polarized proteins to their appropriate domains is a fundamental question in cell biology. The sorting of basolateral proteins is achieved through consensus amino acid motifs serving as molecular zip codes, which recruit specific adaptor complexes. In contrast, apical proteins lack simple sorting motifs, and the mechanisms regulating their sorting and delivery remain poorly understood. Although specific mechanisms have been identified for certain apical proteins, these are not generalizable. In this work, we identified a size asymmetry when comparing the cytoplasmic domains of apical and basolateral membrane proteins. The bias of apical proteins towards smaller cytoplasmic domains led us to question whether the cytoplasmic footprint of polarized proteins has functional relevance as a sorting mechanism. To test this hypothesis, we combined RUSH and FKBP-FRB systems to recruit large cargos to the cytoplasmic domains of apical proteins, to assess the impact on traffic dynamics. We demonstrated that recruitment of large cytoplasmic cargo to apical proteins substantially alters dynamics at the TGN and impairs apical delivery. Moreover, the effect was not dependent on the specific amino acid sequence of the cytoplasmic domains. Interestingly, kinetic impairment did not scale with increased cargo size once a certain threshold was met, suggesting the size filter functions in a binary manner, below which traffic is unimpeded, and above which it is impaired. We could also conclude that the trafficking impairment is not a result of reduced diffusivity on the membrane, as size has negligible effects on 2D membrane diffusion in the absence of a barrier. Consistent with these predictions from literature, we showed by FRAP that mobilities of cargos at the ER were identical whether bulky cytoplasmic domains were attached or not.

Beyond the FKBP-FRB system, we demonstrated that Crb3 traffics biosynthetically with other Crb complex components (Pals1/Patj). However, the Crb3-Pals1 complex dissociates as Crb3 progresses through the Golgi. In accord with our proposed model, a mutant Pals1 which cannot dissociate during Golgi progression significantly impeded the forward trafficking of Crb3. Taken together, these data support an unanticipated mechanism, through which the short cytoplasmic footprint of apical proteins serves as a physical sorting cue (**Fig. 7D**).

At present, the identity and composition of the TGN size filter is unclear. One possibility is that cytoskeletal components make close contact with the TGN at the neck of tubulovesicular budding sites, restricting entry of bulky cargos from diffusing along the membrane into apically destined compartments. Actin and other cytoskeletal components interact with the TGN^27^, including spectrins, which form a tight lattice meshwork along parts of the TGN membrane and play a role in maintaining Golgi structure^28^, but whether they are involved in apical sorting remains to be determined.

## Acknowledgments

We thank Franck Perez, George Church, Fernando Martín-Belmonte, Feng Zhang, Li-En Jao, Didier Trono, and Pantelis Tsoulfas for the kind gift of vectors, and Luke Lavis for the kind gift of JF-Halo and -SNAP dyes. We also thank Macara lab members for valuable comments and advice. Microscopy was performed in part through the Vanderbilt University Cell Imaging Shared Resource. FACS was performed with assistance from the Vanderbilt Technologies for Advanced Genomics (VANTAGE) Flow Cytometry Shared Resource. Some figure diagrams were generated using BioRender. This work was funded by National Institutes of Health grant GM070902.

## Author contributions

CdC proposed the model, designed and built the experimental system, and performed the experiments. IGM supervised the project.

## Materials and Methods

### Reagents

A tabulated list of reagents used in this study are available in Auxiliary Material 2.

### Plasmids and plasmid construction

The following plasmids were received as gifts from the principal investigator listed : pCDH-StrKDEL-neomycin (Franck Perez, Addgene #65307), pCDH-SBP-EGFP-Ecadherin (Franck Perez, Addgene #65290), pCDH-SBP-EGFP-Crb3 (Fernando Martín-Belmonte), lentiCRISPRv2-Puro (Feng Zhang, Addgene #52961), pCMVsp6-nls-hCas9-nls (Li-En Jao), pMD2.G (Didier Trono, Addgene #12259), psPAX2 (Didier Trono, #12260), gRNA-Cloning-Vector (George Church, Addgene #41824), pLV-Golgi eGFP (Pantelis Tsoulfas, Addgene #79809).

All plasmids were grown in Stbl3 bacteria. All plasmid sequences were verified by Sanger sequencing. For PCR involved in cloning, Phusion High-Fidelity DNA Polymerase was used, except in the case of quick-change mutagenesis, where Q5 High-Fidelity Polymerase was used. All oligonucleotides were ordered from IDT. All synthetic gene fragments were ordered as geneBlocks from IDT. All ligations were performed using T4 DNA Ligase. Annotated plasmid maps are available upon request as Snapgene files.

pWPI-StrKDEL was cloned via PCR-amplification of the StrKDEL coding sequence from pCDH-StrKDEL-neomycin using primers to attach a BamHI site at the 5’ end, and a BstBI site at the 3’ end. The PCR product was digested with BamHI and BstBI and ligated into pWPI-MCS-IRES-mScarlet, which was digested with the same eznymes to replace the IRES-mScarlet region.

pWPI-SBP-HaloTag-Crb3 was cloned by inserting a PCR-amplified geneBlock containing the SBP-HaloTag-Crb3 sequence into pWPI-MCS-IRES-mScarlet via BamHI and BstBI digestion to replace the IRES-mScarlet region. pWPI-SBP-HaloTag-Crb3-FKBP was generated by digesting pWPI-SBP-HaloTag-Crb3 with SmaI and BstBI to excise a portion of the HaloTag and Crb3, and replace it with a PCR-amplified geneBlock sequence containing the excised region, a linker, an FKBP sequence, and a stop codon. The insert was flanked by SmaI and BstBI sites to enable digestion and ligation into the cut vector.

pWPI-SNAPtagx1-FRB was generated by inserting a PCR-amplified geneBlock sequence of BglII-SNAPtag-BamHI-FRB-BstBI via BglII + BstBI digestion, and ligating it into a BamHI + BstBI digested pWPI-MCS-IRES-mScarlet vector to replace the IRES-mScarlet region. This process destroys the 5’ BamHI site, but leaves an intact BamHI site between the SNAPtag and FRB. pWPI-SNAPtagx2-FRB was generated by digesting the same geneBlock with BglII and BamHI to isolate the SNAPtag and insert it into a BamHI digested pWPI-SNAPtagx1-FRB. pWPI-SNAPtagx3-FRB was generated the same way by cloning the SNAPtag into pWPI-SNAPtagx2-FRB. pWPI-mCherry-FRB was generated the same way as pWPI-SNAPtagx1-FRB but with mCherry in place of SNAPtag.

pCDH-SBP-EGFP-Crb3-E117STOP was generated by mutating pCDH-SBP-EGFP-Crb3 at the E117 codon from GAG to TAG to create a premature stop codon via site-directed mutagenesis.

pLVX-SBP-mNeonGreen-Ace2 and pLVX-SBP-HaloTag-Ace2-FKBP were designed and ordered from Vector-Builder.

lentiCRISPRv2 : Guide RNAs against mouse *Mpp5* (Pals1) were designed using CHOPCHOP (*27*). MsMpp5-sg1 (Pals1-sg1): GAAAGTATCGGCCATTACGG; MsMpp5-sg2 (Pals1-sg2): TATGAGACGTAGGCGCGAAG; or a nontargeting sequence NT-sg1 : GCGAGGTATTCGGCTCCGCG were cloned into lentiCRISPRv2-Puro as previously described (*28*). Briefly, vector annealing sequences were added to the above guide sequences and their complements and were ordered as oligonucleotides. Then, the base lentiCRISPRv2-Puro vector was digested using BsmBI (EspIII) to excise a 2kbp filler region. Annealed oligonucleotide duplexes were then ligated into the digested vector.

pLVTHM-mApplePals1 was generated by digesting the Pals1 coding sequence from pK-VenusPals1 using BamHI and ligating it into a modified pLVTHM-mApple vector containing an MCS prior to the mApple stop codon. pLVTHM-EGFPPals1ΔN was generated by PCR amplifying the coding sequence corresponding to Pals1 PDZ-SH3-GUK domains (Amino acids Ser245-end) using primers to attach BamHI restriction sites, then digesting the PCR product with BamHI, and ligating it into a modified pLVTHM-EGFP vector containing an MCS prior to the EGFP stop codon.

Knock-in vectors : gCV-mCrb3-sgRNA-KI3 and gCV-mMpp5-sgRNA-KI33B contain guide RNAs used for generating knock-ins. Targeting sequences for Crumbs3 Exon 2 (TGTGAAAGGGTCCGGTGCTG) and Mpp5/Pals1 Exon 3 (ATTCATATATGATGTTGTCA) were identified using CHOPCHOP. Corresponding sequences were cloned into gRNA-Cloning-Vector as described in the protocol on the vector’s Addgene web page, using “Option B” via AflII digestion and Gibson assembly. The insert for Litmus29-SBPHaloCrb3-HDR vector was produced by overlap-extension PCR to attach Crumbs3 homology arms to either side of an SBP-HaloTag sequence, each of which were ordered as geneBlocks. The homology arms each had EcoRV sites near their distal ends to enable EcoRV digestion and ligation into an EcoRV digested Litmus29 vector. The insert for Litmus29-SNAPPals1-HDR vector was designed and ordered as a geneBlock containing Mpp5 homology arms on either side of a SNAPtag sequence, and was cloned into an EcoRV digested Litmus29 by Gibson assembly.

### Cell Culture

Eph4 cells were a gift from Dr. Jurgen Knoblich (Institute of Molecular Biotechnology, Vienna, Austria). HEK293T cells were obtained from the ATCC. EpH4 and HEK293T cells were cultured in DMEM with 4.5g/L D-glucose, L-Glutamine, and 110mg/L sodium pyruvate, supplemented with 10% fetal bovine serum (Gibco), and 1% penicillin/streptomycin (Life Technologies). Cells were grown in humidified incubators supplied with 5% CO2. Cells were passaged at 80-90% confluence using 0.25% trypsin, and reseeded at 1:20.

### CRISPR/Cas9 knock-in generation

To generate endogenously labelled fusion proteins, EpH4 cells were pretreated for 2 days with 5uM farrerol, then one million cells were electroporated in suspension with 550fmol of each pCMV-NLS-hCas9-NLS, a gene-specific gRNA Cloning vector, and an HDR template with 300bp or 850bp homology arms linearized from a Litmus29 vector via EcoRV digestion. (Kit V nucleofection solution, Amaxa program T016). One week following electroporation, cells were labelled with respective JF549-Halo or JF552-SNAPcp dyes, and FACS sorted to generate single cell clones. To confirm HDR knock-in, clones were genotyped via PCR with one primer flanking genomic DNA outside of the insert + homology arms, and one primer inside of the homology arms. Double knock-ins were generated sequentially, by knocking-in a SNAPtag to the *Mpp5* (Pals1) locus in *Crb3*^SBP-Halo/SBP-Halo^ cells.

### Lentiviral production and transduction

Lentiviruses were produced in low-passage HEK293T cells, by calcium phosphate transfection with appropriate lentivectors and lentiviral packaging vectors psPAX2 and pMD2.G. Fresh medium was exchanged 16 hours following transfection. Viral supernatants were collected 48 hrs following transfection, spun down to remove cell debris, and frozen at -80°C. To transduce cells, viral supernatants were added to resuspended EpH4 cells in an Eppendorf tube, and shaken at 400rpm at 37°C for 2 hours prior to plating. To create stable lines, depending on the construct transduced cells were enriched either by selection in puromycin at 1μg/mL for 5 days, or by FACS when the construct was fluorescently sortable. All transductions were performed sparsely to favor single-copy integration, except in the cases of StrKDEL, SNAPtag- or mCherry-FRB transduction where stoichiometric excess relative to RUSH cargoes was required.

### Immunofluorescence

Cells were grown to confluence on Lab-Tek II #1.5 chamber slides, and fixed in 4% paraformaldehyde in pH7.4 PBS for 10 min. After fixation, samples were permeabilized using 0.2% Triton X-100 in PBS for 5 min, before blocking in 10% normal goat serum in PBS for one hour. Primary and secondary antibodies were diluted in blocking buffer at concentrations listed in Supplementary Data 2. Samples were incubated with primary and secondary antibodies sequentially and washed 4x in PBS for 5 min. following each incubation.

### Confocal Imaging

Confocal images and videos were acquired using a Nikon A1-R at 1024×1024 resolution using either the Galvano or resonant scanner, as indicated. Either a 40× / 1.3NA, 60× / 1.4NA, or 100× / 1.4NA Plan Apochromat objective was used. For visualization, XZ orthogonal views have been denoised using Nikon’s Denoise.ai package. No quantitative data were extracted from denoised images.

### Near-TIRF Imaging

Near-TIRF images and videos were acquired using a Nikon TIRF equipped with LUN-F multi-excitation diode laser lines and Apo-TIRF 100× / 1.49NA oil immersion lens with a Photometrics Prime 95B sCMOS camera for detection. The 561nm laser line was used to excite JF552-SNAPcp, or JF549-Halo dyes. The 641nm laser line was used to excite JF646-Halo or JFX646-Halo dyes. When imaging both channels, lasers were fired simultaneously. Images were captured on the camera simultaneously by use of an Optosplit III. To enter near-TIRF, laser angles were manually adjusted with the aid of a Bertrand lens camera to visualize position of the lasers on the rear of the objective. Live imaging was performed in a stage-top incubator (Tokai) heated to 37°C. Videos were acquired at 10Hz.

### RUSH Assays

RUSH assays were performed using EpH4 cell lines containing endoplasmic reticulum retention hooks in the form of lentiviral Streptavidin-KDEL, along with cargo fusion proteins containing a streptavidin-binding peptide and fluorescent reporter. EpH4 RUSH cells were grown to confluence on Lab-Tek II #1.5 chambered coverglass. For live cell imaging experiments, slides were placed within a stage-top incubator (Tokai) mounted on a Nikon A1R, set at 37oC and supplied with 5% CO2. To activate the RUSH system, biotin was added to each well at a final concentration of 80μM.

### FKBP-FRB Dimerization

For experiments using the FKBP-FRB dimerization system, cells were pre-treated 1 hour prior to the assay with A/C Heterodimerizer (Takara) diluted at 1:200 in complete media concurrent with JF-Halo or -SNAP dye labeling. A/C Heterodimerizer was also included at the same concentration in the media during the experiment.

### Fluorescence recovery after photobleaching (FRAP)

Cells were grown to confluence on Lab-Tek II #1.5 chambered coverglass. For FRAP experiments, slides were placed within a stage-top incubator mounted on a Nikon A1R, set at 37oC and supplied with 5% CO2. To assess relative diffusivity of target proteins at the ER, biotin was withheld from culture media so that Crb3 RUSH cargo was retained there. Using this signal, we focused on a medial portion of cell where the reticular pattern of RUSH cargo was observed to demarcate the ER. A circular RoI was created for the FRAP region. Prior to photobleaching, several frames were acquired as a reference for initial fluorescence intensity. After which, the FRAP RoI was stimulated with lasers corresponding to the fluorophores being bleached, for either 4 seconds (in Fig. 3H, I) or 2 seconds (in Fig. 5B). Immediately after stimulation, recovery frames were acquired every 3 seconds for 3 minutes (in Fig. 3H, I), or every 0.5 seconds for 2 minutes (in Fig. 5B). Following acquisition, a reference RoI was drawn within the same cell distal to the stimulation RoI to account for background photobleaching during acquisition. Reference corrected time series data of the photobleached RoI were exported from Nikon Elements. These data were then imported into R, where intensity values were maximum normalized to 1, per channel and per cell. Half-maximum recovery times were determined based on the time at which normalized intensity recovered halfway between the intensity at photobleaching, and the plateau of recovery.

### SDS-Page and Immunoblotting

Cells were washed with ice-cold PBS prior to on-plate cell lysis in RIPA buffer (10 mM Tris-HCl pH 8.0, 1 mM EDTA, 0.5 mM EGTA, 1% Triton X-100, 0.1% sodium deoxycholate, 0.1% SDS, 140 mM NaCl) supplemented with 1x protease and phosphatase inhibitor cocktail. Lysates were collected and incubated on ice for 15 min. before centrifugation at 18,000g for 15 min. at 4°C. Supernatants were then collected and protein concentration was determined by Precision Red spectrophotometry on a NanoDrop® ND-1000. For each sample, 35μg protein was prepared in Laemli sample buffer w/ 2-Mercaptoethanol, boiled at 98°C for 5 min., then loaded and resolved via SDS-PAGE on a 4% stacking/10% resolving bis-acrylamide gel. Resolved proteins were wet transferred onto a PVDF membrane at 80mA for 16 hours at 4°C.

Membranes were blocked in 1x Western blocking reagent in TBST for 1 hour at room temperature, prior to overnight incubation at 4°C with primary antibodies (Rabbit anti-Pals1, 1:500; or Mouse anti-β-Actin, 1:2000) diluted in blocking buffer on a shaker. Membranes were washed 4x for 10 min on a shaker in TBST (10 mM Tris–HCl pH 8, 150 mM NaCl, 0.05% Tween 20). Membranes were incubated with HRP-conjugated secondary antibodies (goat anti-rabbit 1:1000; or goat anti-mouse 1:5000) diluted in blocking reagent for 1 hour at room temperature on a shaker. Membranes were washed 4x for 10 min on a shaker in TBST, before development in chemiluminescent substrate. Images of developing membranes were acquired on an Amersham Imager 600. Quantification of band intensity was performed in ImageJ, by drawing a box of uniform size around each band and extracting mean intensity data. Background correction was performed by subtracting the mean intensity of the adjacent membrane.

## Quantification and Statistics

### RUSH quantification

To quantify trafficking dynamics during RUSH experiments, we extracted the fluorescence intensity over time at the presumptive Golgi apparatus for each cell. To do this in Nikon Elements, we generated maximum intensity projections in the Z-plane, followed by a maximum intensity projection in time to identify bright regions of interest (RoI) corresponding to the Golgi apparatus. Oversaturated RoIs were excluded from analysis as their maximum intensity would underestimated. Using these RoIs, we extracted mean fluorescence intensity values for the Golgi of each cell, for each channel, at each time point in the experiment. These values represent the raw intensity data, *I*_*raw*_. To compare intensity information between channels and cells, we rescaled *I*_*raw*_ for each channel and cell via min-max normalization on a (0,1) scale, such that for any time *t*:

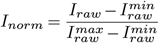

Where 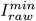 and 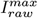 are the per-series *I*_*raw*_ minimum and maximum respectively. Using these normalized data, we extracted temporal information related to changes in intensity by identifying the times during the experiment at which certain *I*_*norm*_ values were reached for each channel and cell. *t*_*peak*_ is the time at which *I*_*norm*_ reached maximum (1), corresponding to the time at which the maximum amount of RUSH cargo was present at the Golgi. *t*_*prior*_ and *t*_*post*_ are the times at which *I*_*norm*_ reached half-maximum value (0.5) on either side of the peak, with *t*_*prior*_ corresponding to the time half the RUSH cargo accumulated at the Golgi, and *t*_*post*_ corresponding to the time half the RUSH cargo became depleted from the Golgi. Half-maximum quantiles were chosen because the curves appear linear at this portion, and to minimize effects of noise which occur by using quantiles near inflection points:

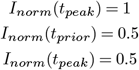

To account for slight variation in time to activation of the RUSH system between experiments, we used differences between these *t* values to generate time values between relative positions on the curve:

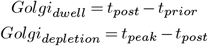

These calculations were made per channel and per cell, then aggregated and plotted. See **Fig. S4A** for visual explanation.

### RUSH Heatmap Visualization

To visualize per-cell dynamics of RUSH experiments in a concise manner, we generated heatmaps using the ComplexHeatmap R package (*29*). The input for heatmaps were matrices of time-interpolated *I*_*norm*_ data, where each row is an individual cell, each column is a time point, with *I*_*norm*_ as element values. Note that heatmaps in figures have been rotated, so rows and columns are flipped. For visualization purposes, cells are arranged in descending order based on their *t*_*post*_ value. When multiple channels are being visualized, the ordering is based on a single channel, but both heatmaps will use the same order so that traces within the same cell can be compared. For coloring, an inverted ‘RdYlBu’ palette was used: with blue, white, and red colors representing minimum, middle, and maximum values respectively. See **Fig. S4B** for visual explanation.

### Plot graphics

All figure plots were created in R using ggplot2 included in the tidyverse package. Where applicable, graphs are formatted as SuperPlots, where individual data points are plotted and colored by experiment, along with the per-experiments means as larger diamonds (*30*).

### Statistics

Statistical tests were performed as indicated in figure legends, using base R functions. To determine whether to use a parametric or nonparametric test, homoscedasticity was assessed by Levene’s test, and cases with Levene’s p < 0.05 were considered to have unequal variance prescribing use of a nonparametric test. All statistical tests were performed on experimental means, except in FRAP experiments where statistics were performed on per-cell recovery values, though experimental means are also plotted. In plot graphics: Error bars represent the mean and standard deviation of experimental means. Statistics are summarized using the following indicators: *n*.*s*. : *p >* 0.05. * : *p <* 0.05. ** : *p <* 0.01. *** : *p <* 0.001.

## Supplementary Figures

**Figure S1.**
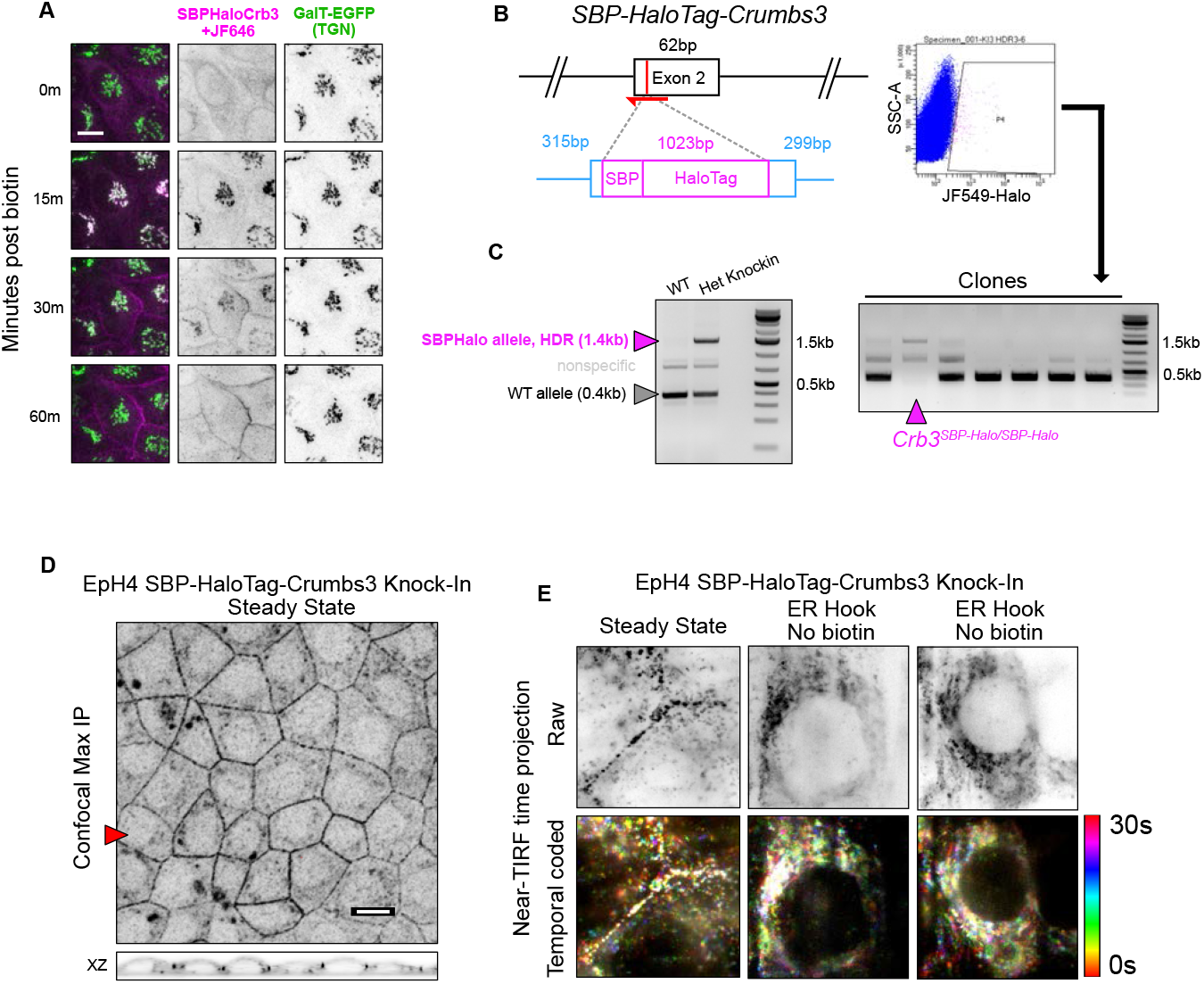
**(A)** Confocal imaging of exogenous SBP-HaloTag-Crb3 RUSH in cells also expressing trans-Golgi reporter GalT-EGFP. Confocal maximum intensity projections. Scalebar 10μm. **(B)** Left: Schematic of CRISPR/Cas9 gene editing strategy to insert the *SBP-HaloTag* sequence into *Crb3* exon 2, which encodes amino acids immediately following the signal peptide of Crb3. In red: guide RNA targeting and approximate cut site. In magenta: *SBP-HaloTag* insert. In blue: homology arms flanking the insert to facilitate homology directed repair (HDR). Right: FACS gating to isolate JF549-Halo+ clones. **(C)** Genotyping for SBP-Halo HDR alleles in putative knock-in clones. PCR product lengths: wild-type allele: 385bp; HDR allele: 1408bp; nonhomologous end-joining allele: 1700 and 2100bp (none shown). **(D)** Confocal microscopy of EpH4 *Crb3SBP-Halo/SBP-Halo* cells at steady state labeled with JF549-Halo. Maximum intensity projection or XZ orthogonal view. Scalebar 10μm. 100×100μm. **(E)** Near-TIRF of EpH4 *Crb3*^*SBP-Halo/SBP-Halo*^ cells at steady state labeled with JF549-Halo and imaged live, at either steady state (junctional) or with the StrKDEL hook added (ER localized). Images are maximum intensity projections in time over a 30 second span. Top row: raw data. Bottom row: temporal color coded.

**Figure S2.**
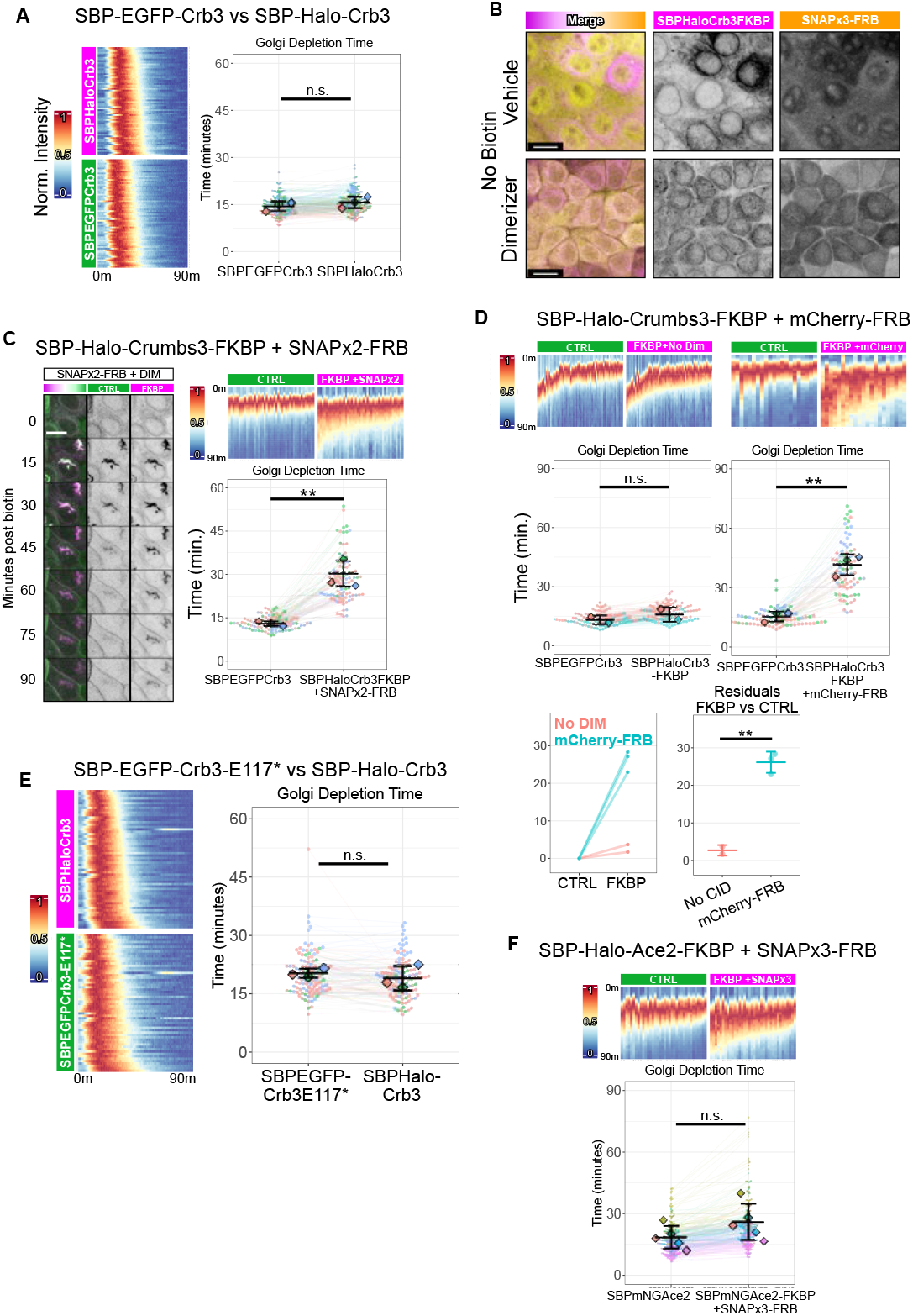
**(A)** Golgi trafficking dynamics of SBP-EGFP-Crb3 versus SBP-Halo-Crb3. n = 179 total cells. N = 3 experiments. Student’s t-test. n.s. p = 0.398. **(B)** Visualization of SNAPtagx3-FRB in cells expressing ER-retained SBP-Halo-Crb3-FKBP, with either vehicle or with CID. Note nuclear localization in vehicle, which transitions to colocalize with the FKBP-fused construct following addition of CID. **(C)** Extended data related to Fig. 3D-G, using the same methodology but with SNAPtagx2-FRB as recruitable cargo in the presence of CID. n = 90 total cells. N = 3 experiments. Student’s t-test. **p = 0.00256. **(D)** Extended data related to Fig. 3D-G, using the same methodology but with mCherry-FRB as recruitable cargo, and either vehicle or CID. Vehicle: n = 94 total cells. N = 2 experiments. Welch’s t-test. n.s. p = 0.481. CID: n = 74 total cells. N = 3 experiments. Student’s t-test. **p = 0.00155. Residuals: One-way ANOVA. **p = 0.00184. **(E)** Golgi trafficking dynamics of mutant SBP-EGFP-Crb3-E117STOP, lacking the PDZ-binding motif, versus full length SBP-Halo-Crb3. n = 108 total cells. N = 3 experiments. Student’s t-test. n.s. p = 0.547. **(F)** Extended data related to Fig. 4C-D, using the same methodology but with SNAPtagx3-FRB as recruitable cargo in the presence of CID. n = 228 total cells. N = 5 experiments. Student’s t-test. n.s. p = 0.145.

**Figure S3.**
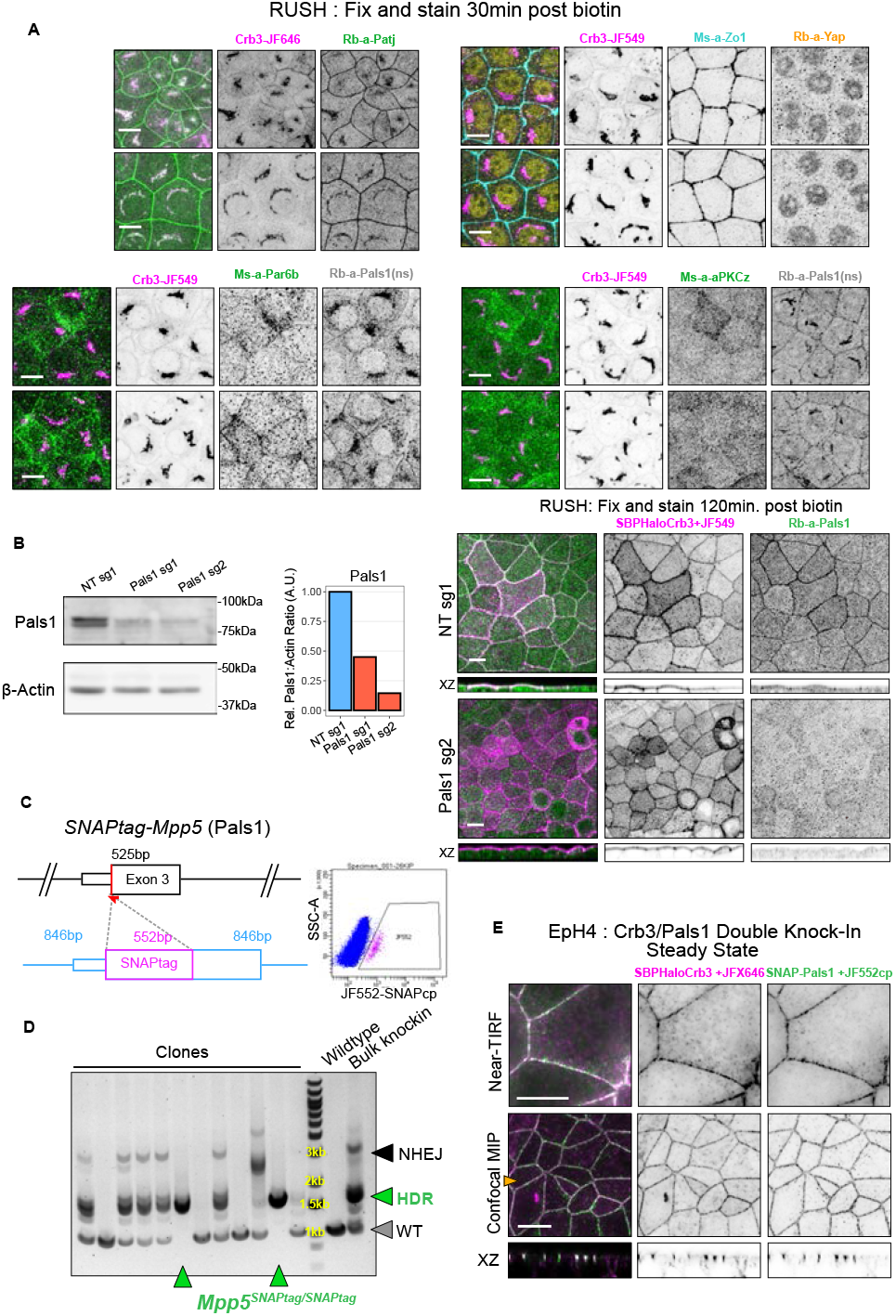
**(A)** Images of SBP-HaloTag-Crb3 RUSH cells fixed after 30 minutes biotin, and immunostained against endogenous protein targets as indicated. Confocal maximum intensity projections. **(B)** Left: Immunoblot of SBP-HaloTag-Crb3 RUSH cells transduced with pLentiCRISPRv2 virus with nontargeting (NT), or Pals1 sgRNAs. Right: Images of nontargeting, or Pals1 deleted SBP-HaloTag-Crb3 RUSH cells fixed after 120 minutes biotin, and immunostained against endogenous Pals1. **(C)** Left: Schematic of CRISPR/Cas9 gene editing strategy to insert the SNAPtag sequence into *Mpp5* (Pals1) exon 3, the first coding exon of the gene. In red: guide RNA targeting and approximate cut site. In magenta: *SNAPtag* insert. In blue: homology arms flanking the insert to facilitate homology directed repair (HDR). Right: FACS gating to isolate JF552-SNAPcp+ clones. **(D)** Genotyping for *SNAPtag* HDR alleles in putative knock-in clones. PCR product lengths: wild-type allele: 1054bp; HDR allele: 1606bp; nonhomologous end-joining allele: 3300 and 2400bp. **(E)** Confocal microscopy of EpH4 *Crb3SBP-Halo/SBP-Halo Mpp5SNAPtag/SNAPtag* double knock-in cells at steady state labeled with JFX646-Halo and JF552-SNAPcp.

**Figure S4.**
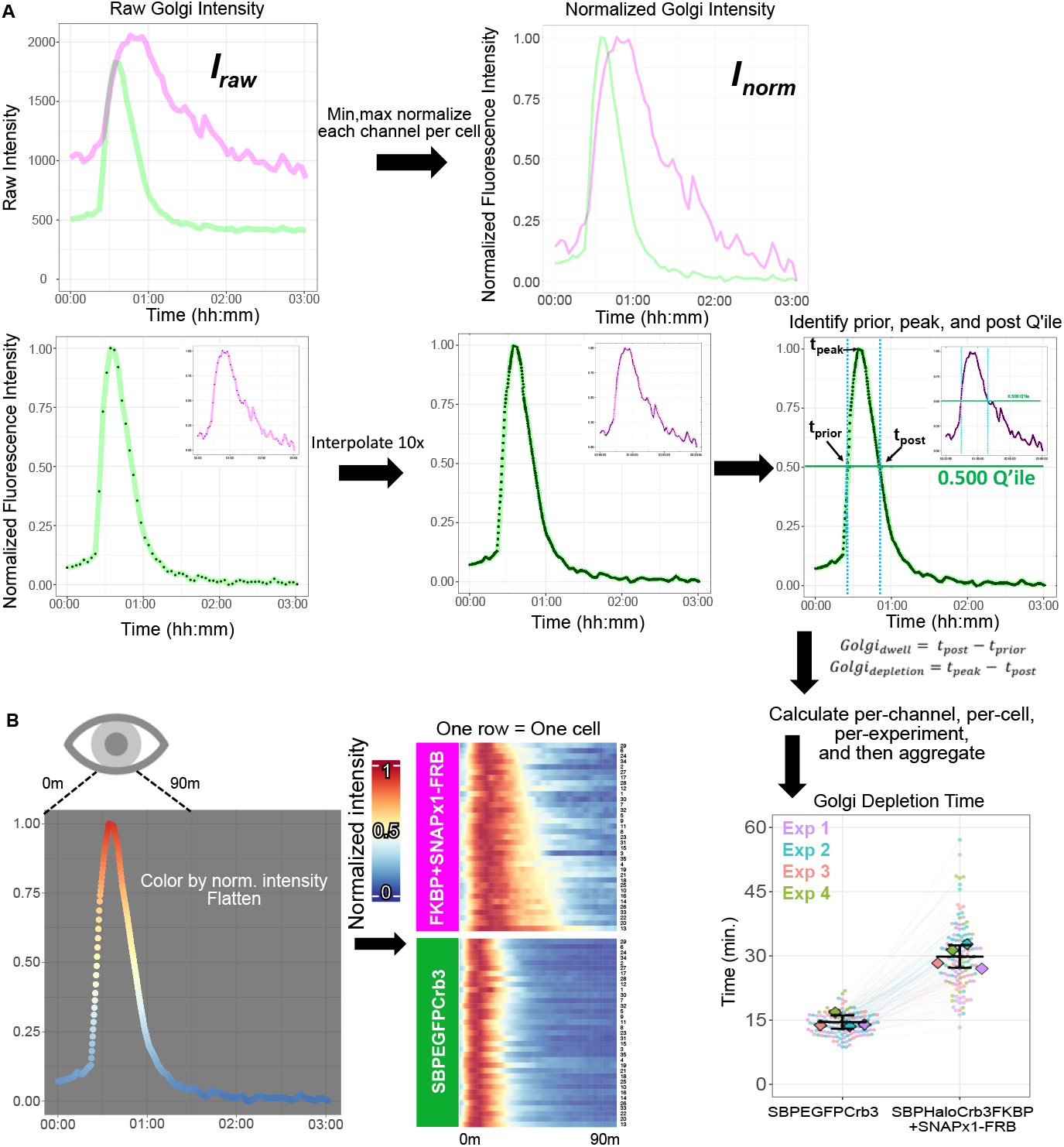
**(A)** RUSH normalization and quantification strategy, as detailed in Methods: “RUSH quantification” **(B)** RUSH heatmap visualization, as detailed in Methods: “RUSH Heatmap Visualization”

## Supplementary Videos

**Movie S1**.

Related to Fig. 2C: RUSH of exogenous SBP-HaloTag-Crumbs3 in polarized EpH4 cells. Top: Confocal maximum intensity projection. Bottom: Denoised XZ orthogonal view. Scalebar: 10μm. Time formatting: hh:mm:ss.

**Movie S2**.

Related to Fig. 2D: RUSH of endogenous SBP-HaloTag-Crumbs3 in polarized EpH4 cells. Confocal Z-slice. Scale-bar: 10μm. Time formatting: hh:mm:ss.

**Movie S3**.

Related to Fig. 2E: RUSH of exogenous SBP-HaloTag-Crumbs3 and SBP-EGFP-Ecadherin in polarized EpH4 cells. Confocal denoised XZ orthogonal view. Scalebar 10μm. Time formatting: hh:mm:ss.

**Movie S4**.

Related to Fig. 3D: RUSH of exogenous SBP-EGFP-Crumbs3 and SBP-HaloTag-Crumbs3-FKBP in polarized EpH4 cells. Top: with SNAPx1-FRB and vehicle. Middle: With SNAPx1-FRB and CID. Bottom: With SNAPx3-FRB and CID. Confocal maximum intensity projection Scalebar 10μm. Time formatting: hh:mm:ss.

**Movie S5**.

Related to Fig. 4C: RUSH of exogenous SBP-mNeonGreen-Ace2 and SBP-HaloTag-Ace2-FKBP in polarized EpH4 cells. Top: with SNAPx1-FRB and vehicle. Middle: With SNAPx1-FRB and CID. Bottom: With SNAPx3-FRB and CID. Confocal maximum intensity projection Scalebar 10μm. Time formatting: hh:mm:ss.

**Movie S6**.

Related to Fig. 5C: RUSH of exogenous SBP-HaloTag-Crumbs3 in polarized EpH4 cells also expressing mApple-Pals1. Confocal maximum intensity projection Scalebar 10μm. Time formatting: hh:mm:ss.

**Movie S7**.

Related to Fig. 5D: RUSH of exogenous of exogenous SBP-HaloTag-Crumbs3 in polarized EpH4 cells also expressing mApple-Pals1. Note evolution of Pals1 positive, Crumbs3 negative puncta around 17:45. Denoised 3D Alpha projection. Bounding box is 60μm x 60μm. Time formatting: hh:mm:ss.

**Auxiliary Material 1**.

Related to Fig 1: List of polarized membrane proteins from Zeke et al., 2021 detailing their localization, cytoplasmic and total amino acid lengths.

**Auxiliary Material 2**.

Spreadsheet of vector constructs, dyes, reagents, and antibodies used in this study, along with catalog numbers where applicable.

## References

1. E. Rodriguez-Boulan, I. G. Macara, Organization and execution of the epithelial polarity programme. Nat Rev Mol Cell Biol 15, 225–242 (2014).

2. E. H. Stoops, M. J. Caplan, Trafficking to the apical and basolateral membranes in polarized epithelial cells. J Am Soc Nephrol 25, 1375–1386 (2014).

3. R. Legouis et al., Basolateral targeting by leucine-rich repeat domains in epithelial cells. EMBO Rep 4, 1096–1102 (2003).

4. K. C. Miranda et al., A dileucine motif targets E-cadherin to the basolateral cell surface in Madin-Darby canine kidney and LLC-PK1 epithelial cells. J Biol Chem 276, 22565–22572 (2001).

5. R. T. Youker et al., Multiple motifs regulate apical sorting of p75 via a mechanism that involves dimerization and higher-order oligomerization. Mol Biol Cell 24, 1996–2007 (2013).

6. P. Scheiffele, J. Peranen, K. Simons, N-glycans as apical sorting signals in epithelial cells. Nature 378, 96–98 (1995).

7. D. S. Levic, M. Bagnat, Self-organization of apical membrane protein sorting in epithelial cells. FEBS J 289, 659–670 (2022).

8. D. S. Levic et al., Distinct roles for luminal acidification in apical protein sorting and trafficking in zebrafish. J Cell Biol 219, (2020).

9. C. L. Kinlough et al., Core-glycosylated mucin-like repeats from MUC1 are an apical targeting signal. J Biol Chem 286, 39072–39081 (2011).

10. S. Paladino, T. Pocard, M. A. Catino, C. Zurzolo, GPI-anchored proteins are directly targeted to the apical surface in fully polarized MDCK cells. J Cell Biol 172, 1023–1034 (2006).

11. O. Makarova, M. H. Roh, C. J. Liu, S. Laurinec, B. Margolis, Mammalian Crumbs3 is a small transmem-brane protein linked to protein associated with Lin-7 (Pals1). Gene 302, 21–29 (2003).

12. Y. T. Chen, D. B. Stewart, W. J. Nelson, Coupling assembly of the E-cadherin/beta-catenin complex to efficient endoplasmic reticulum exit and basal-lateral membrane targeting of E-cadherin in polarized MDCK cells. J Cell Biol 144, 687–699 (1999).

13. J. Schlessinger, Cell signaling by receptor tyrosine kinases. Cell 103, 211–225 (2000).

14. C. P. Lusk, G. Blobel, M. C. King, Highway to the inner nuclear membrane: rules for the road. Nat Rev Mol Cell Biol 8, 414–420 (2007).

15. R. Ungricht, M. Klann, P. Horvath, U. Kutay, Diffusion and retention are major determinants of protein targeting to the inner nuclear membrane. J Cell Biol 209, 687–703 (2015).

16. D. K. Breslow, E. F. Koslover, F. Seydel, A. J. Spakowitz, M. V. Nachury, An in vitro assay for entry into cilia reveals unique properties of the soluble diffusion barrier. J Cell Biol 203, 129–147 (2013).

17. G. Boncompain et al., Synchronization of secretory protein traffic in populations of cells. Nat Methods 9, 493–498 (2012).

18. M. H. Roh et al., The Maguk protein, Pals1, functions as an adapter, linking mammalian homologues of Crumbs and Discs Lost. J Cell Biol 157, 161–172 (2002).

19. T. W. Hurd, L. Gao, M. H. Roh, I. G. Macara, B. Margolis, Direct interaction of two polarity complexes implicated in epithelial tight junction assembly. Nat Cell Biol 5, 137–142 (2003).

20. A. Zeke, L. Dobson, L. I. Szekeres, T. Lango, G. E. Tusnady, PolarProtDb: A Database of Transmembrane and Secreted Proteins showing Apical-Basal Polarity. J Mol Biol 433, 166705 (2021).

21. Y. H. Lin et al., AP-2-complex-mediated endocytosis of Drosophila Crumbs regulates polarity by antagonizing Stardust. J Cell Sci 128, 4538–4549 (2015).

22. J. R. Houser et al., The impact of physiological crowding on the diffusivity of membrane bound proteins. Soft Matter 12, 2127–2134 (2016).

23. K. Weiss et al., Quantifying the diffusion of membrane proteins and peptides in black lipid membranes with 2-focus fluorescence correlation spectroscopy. Biophys J 105, 455–462 (2013).

24. F. Rouaud, I. Mean, S. Citi, The ACE2 Receptor for Coronavirus Entry Is Localized at Apical Cell-Cell Junctions of Epithelial Cells. Cells 11, (2022).

25. G. Egea, C. Serra-Peinado, M. P. Gavilan, R. M. Rios, Cytoskeleton and Golgi-apparatus interactions: a two-way road of function and structure. Cell Health and Cytoskeleton, (2015).

26. K. A. Beck, Spectrins and the Golgi. Biochim Biophys Acta 1744, 374–382 (2005).

27. K. Labun et al., CHOPCHOP v3: expanding the CRISPR web toolbox beyond genome editing. Nucleic Acids Res 47, W171–W174 (2019).

28. N. E. Sanjana, O. Shalem, F. Zhang, Improved vectors and genome-wide libraries for CRISPR screening. Nature Methods 11, 783–784 (2014).

29. Z. Gu, R. Eils, M. Schlesner, Complex heatmaps reveal patterns and correlations in multidimensional genomic data. Bioinformatics 32, 2847–2849 (2016).

30. S. J. Lord, K. B. Velle, R. D. Mullins, L. K. Fritz-Laylin, SuperPlots: Communicating reproducibility and variability in cell biology. J Cell Biol 219, (2020).

